# Adeno-associated virus (AAV) reduces cortical dendritic complexity in a TLR9-dependent manner

**DOI:** 10.1101/2021.09.28.462148

**Authors:** Christos M. Suriano, Jessica L. Verpeut, Neerav Kumar, Jie Ma, Caroline Jung, Lisa M. Boulanger

## Abstract

Recombinant adeno-associated viruses (AAVs) allow rapid and efficient gene delivery in the nervous system. AAVs are widely used in research and are the basis of multiple FDA-approved gene therapies. Here, we find that the immune response to AAV’s genome reduces dendritic complexity in mammalian cortex. Dendritic loss associated with AAV-mediated gene delivery occurs at experimentally-relevant titers, cannot be explained by responses to transgene expression or surgery, and is not restricted to a particular capsid serotype, encoded transgene, promoter, or production facility. AAV-associated dendritic loss is accompanied by a decrease in the frequency and amplitude of miniature excitatory postsynaptic currents (mEPSCs) and upregulation of immune molecules that can limit dendritic complexity and synaptic transmission. Blocking detection of unmethylated CpG-rich DNA via Toll-like receptor 9 (TLR9) protects dendritic complexity, suggesting that immunodetection of a core feature of the AAV genome triggers dendritic loss. These results reveal previously unsuspected impacts of AAV on neuronal structure and function and identify TLR9 inhibitors as important tools to improve the safety and efficacy of AAV-mediated gene delivery in the nervous system.

## Introduction

Clinical and basic neuroscience have been revolutionized by recombinant viral technology. Recombinant adeno-associated viruses (AAVs) are among the most commonly used viruses in the central nervous system (CNS), because they are non-pathogenic, non-replicative, have broad tropism, and drive strong gene expression in postmitotic neurons ^1^. In neuroscience research, AAV is widely used to express exogenous proteins, including circuit tracers, calcium indicators, and optogenetic ion channels for research purposes. Clinically, AAV is used to restore endogenous protein function to treat Leber’s congenital amaurosis 2 ^2,3^ and spinal muscular atrophy ^4,5^. Additional AAV-based clinical trials are underway to treat a number of brain disorders, including amyotrophic lateral sclerosis and Parkinson’s, Canavan’s, and Alzheimer’s diseases.

In the periphery, AAV-mediated gene delivery triggers cellular and humoral immune responses, including upregulation of major histocompatibility complex class I (MHCI) and MHCII proteins, activation of CD8^+^ cytotoxic T cells and CD4^+^ helper T cells, and engagement of B cell mediated antibody responses ^6–10^. Previous exposure to AAV, either due to infection with naturally occurring AAV or administration of engineered AAV, can leave neutralizing antibodies and memory T cells that rapidly clear the virus, limiting transgene expression. In the immune privileged context of the CNS however, the AAV genome persists as a stable episome, allowing sustained transgene expression. The relative stability of transgene expression has contributed to the rapid expansion of AAV-mediated gene delivery in the nervous system.

Even though AAV is not cleared by the immune response in the CNS, neuroinflammation is emerging as an important topic in AAV-mediated gene delivery. CNS-directed AAV delivery has been associated with an increase in neutralizing antibodies ^11–13^, circulating CD45^+^ leukocytes, natural killer cells, and CD8^+^ and CD4^+^ T lymphocytes ^14–16^. Elements of the AAV genome, specifically, have been associated with toxicity in the retina and hippocampus ^17–19^. AAV’s genome is detected by Toll-like receptor 9 (TLR9), an innate immunoreceptor that detects unmethylated CpG dinucleotides, a hallmark of microbial DNA ^6–8,20^. Limiting detection of CpG DNA by blocking TLR9 can prevent or delay retinal inflammation and toxicity associated with AAV-mediated gene delivery ^19^. The *TLR9* gene is expressed by microglia, astrocytes, and neurons in the CNS, and responses to CpG-rich DNA have been detected in both astrocytes and microglia ^21–23^. In the periphery, TLR9 activation triggers a multi-faceted immune response, including the upregulation of pro-inflammatory cytokines and components of the complement cascade, MHCI- and MHCII-based antigen presentation, engagement of CD8^+^ T cells and the production of neutralizing antibodies, paralleling changes seen after AAV-mediated gene delivery in the CNS ^6–10,24–27^.

Notably, several immune proteins that are engaged downstream of TLR9 activation have additional roles in the healthy CNS, where they regulate neuronal development, morphology, and synaptic transmission. For example, proteins of the complement cascade (including C1q, C3, and C4) and members of the major histocompatibility complex class I (MHCI; including H2-K and H2-D) contribute to synapse elimination in the developing visual system ^28–33^. H2-K and H2-D also negatively regulate dendritic complexity and the frequency and amplitude of miniature excitatory postsynaptic currents (mEPSCs) in hippocampal and cortical neurons, and limit the levels of Ca^2+^-permeable α-amino-3-hydroxy-5-methyl-4-isoxazolepropionic acid (AMPA) receptors in the lateral geniculate nucleus ^29,30,34–36^. The tyrosine kinase Fyn regulates cortical and hippocampal spine density and morphology ^37^, and the CXCL10 cytokine receptor, CXCR3, negatively regulates neurite length after injury ^38^. The dual roles of these and other immune proteins in neuronal homeostasis raises the possibility that an immune response to AAV-mediated gene delivery could disrupt neuronal structure or function.

Here, we report that experimentally-relevant titers of AAV reduce dendritic complexity and the frequency and amplitude of mEPSCs recorded from pyramidal neurons in somatosensory cortex, three weeks post-injection. Dendritic loss is dose-dependent, is not a response to transgene expression, and is not limited to a particular capsid serotype, encoded transgene, or promoter. AAV-induced dendritic loss is associated with the persistent upregulation of several immune molecules that can negatively regulate dendritic complexity and synaptic transmission. Dendritic simplification is prevented by administering a TLR9 antagonist, suggesting that dendrites are impacted as a consequence of an immune response to unmethylated CpG-rich DNA, a core feature of AAV genomes.

## Results

### AAV-mediated gene delivery reduces dendritic complexity in cortical pyramidal neurons

To determine if AAV-mediated gene delivery affects neuronal morphology, we used DiOlistic labeling to visualize individual pyramidal neurons from primary somatosensory cortex (S1) in injected ipsilateral and non-injected contralateral hemispheres. Unless otherwise noted, measurements were made three weeks post-injection, a common starting point for AAV-based studies. Unilateral injection of 4×10^10^ viral genomes (vg) of AAV8-*hSyn-mCherry* drives *mCherry* expression in S1 of wild type C57BL/6J mice **(Fig. 1a & 3a, Supplemental Fig. 1a & 3a)**, demonstrating successful viral transduction. Cell density does not differ in ipsilateral versus contralateral hemispheres, suggesting that AAV-mediated gene delivery is not associated with significant cell death in cortex **(Supplemental Fig. 1b)**. Dendritic length and Sholl intersections from neurons in the contralateral hemisphere are indistinguishable from untreated controls for all conditions **(Supplemental Fig. 1c-f)**, and therefore morphological data are represented as an ipsilateral/contralateral ratio within animal. Representative tracings of ipsilateral pyramidal neurons are shown in **Fig. 1b**. Following administration of AAV8-*hSyn-mCherry*, total dendritic length and Sholl intersections are significantly reduced in the ipsilateral hemisphere relative to both untreated controls and vehicle-injected animals **(Fig. 1c-d)**. In contrast, dendritic length and Sholl intersections are indistinguishable from untreated controls in vehicle-injected animals, suggesting that dendritic simplification is not a consequence of surgery or injection, but rather reflects a specific response to AAV-mediated gene delivery.

**Figure 1.**
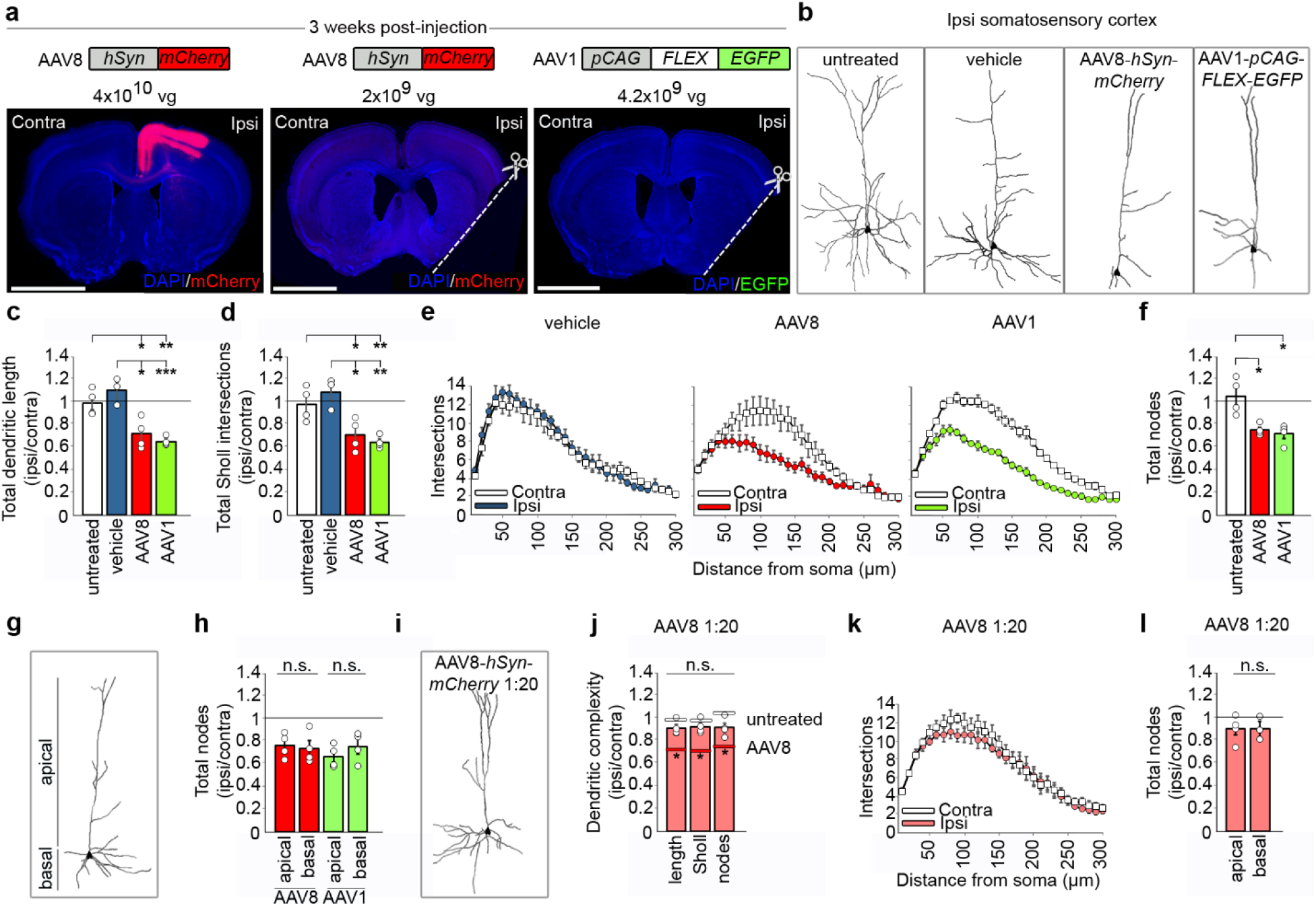
AAV reduces dendritic complexity in pyramidal cells of somatosensory cortex three weeks post-injection. **(a)** AAV8-*hSyn-mCherry* (4×10^10^ vg or 2×10^9^ vg), or AAV1-*pCAG-FLEX-EGFP* (4.2×10^9^ vg) in the absence of CRE, was unilaterally injected into S1. Only the higher dose of AAV8-*hSyn-mCherry* produces detectable transgene expression (red; see also Figure 3a and Supplemental Figure 1a). In cases where transgene expression was not present, the ipsilateral hemisphere was marked in fixed sections by making a ventrolateral incision (dashed line). Blue, DAPI cell body counterstain. Scale bars, 2.5 mm. **(b)** Example tracings of DiO-labeled cells from control or injected ipsilateral hemispheres. **(c-d)** Effects of AAV8 (red) and AAV1 (green) on total dendritic length and Sholl intersections relative to untreated (white) and vehicle (blue) controls. **(e)** Sholl intersections versus distance from the soma. **(f)** Effects of AAV8 and AAV1 on total nodes relative to untreated controls. **(g)** Example trace showing separation of apical and basal compartments. **(h)** Effects of AAV8 and AAV1 on nodes in the apical and basal dendritic compartments. **(i)** Example tracing of DiO labeled cell after AAV8 at 1:20 dilution (2×10^9^ vg). **(j)** Effects of AAV8 1:20 dilution on total dendritic length, Sholl intersections and total nodes relative to untreated controls and full AAV8 dose. **(k)** Effects of AAV8 1:20 dilution on Sholl intersections over distance from soma. **(l)** Effects of AAV8 1:20 dilution on nodes in apical and basal compartments. *n*=4 mice for untreated control, *n*=3 for vehicle, *n*=4 for each AAV8 dose and AAV1; 10-19 neurons per animal (average = 14 cells). Data represented as mean ± S.E.M. *p<0.05; **p<0.01; ***p<0.001.

The effects of AAV8-*hSyn-mCherry* on dendrites could arise from a number of factors, including expression of *mCherry*, an exogenous gene with the potential for neurotoxicity ^39^, or host responses to the viral capsid, genome, promoter, or preparation. To test the importance of transgene expression and other features of this virus in dendritic loss, we unilaterally injected 4.2×10^9^ vg of AAV1-*pCAG-FLEX-EGFP* into S1 of wild type, non-*CRE* expressing C57BL/6J mice. Compared to the AAV used above, this virus has a different capsid serotype (AAV1 vs AAV8), promoter (*pCAG* vs *hSyn*), production facility (Penn Vector Core vs. Princeton Viral Vector Core), and encoded transgene (*EGFP* vs *mCherry*). In addition, the LoxP sites in the *FLEX* plasmid require CRE recombinase for efficient recombination, and therefore this virus should not drive significant *EGFP* expression in the absence of CRE. A low level of CRE-independent recombination can occur during DNA amplification and viral vector production (~0.01-0.03% of viral vectors for a typical production protocol), and therefore we searched for *EGFP* expression in injected hemispheres. EGFP could not be detected by confocal fluorescence microscopy **(Fig. 1a & Supplemental Fig. 1a)** or using immunoblotting to amplify EGFP signals **(Fig. 3a)**. Despite the lack of detectable transgene expression, AAV1-*pCAG-FLEX-EGFP* causes a reduction in total dendritic length and Sholl intersections that closely matches what is seen after AAV8-mediated delivery of *mCherry* **(Fig. 1c-d)**. The reduction in Sholl intersections in both cases is broadly distributed throughout the dendritic arbor **(Fig. 1e)**. These results reveal that AAV-mediated gene delivery causes striking dendritic loss. Furthermore, dendritic loss does not require detectable transgene expression, and is not specific to a particular capsid serotype, promoter, encoded transgene, or production facility, but is triggered by divergent AAV-based gene delivery systems.

The reduction in dendritic length and Sholl intersections by AAV8-*hSyn-mCherry* and AAV1-*pCAG-FLEX-EGFP* could reflect either shortening or elimination of dendritic segments. To discriminate between these possibilities, we counted the total number of nodes (branch points) per neuron. Nodes are lost when entire dendritic segments are eliminated, but not when segments only decrease in length. After AAV injection, the number of nodes in cells of the non-injected contralateral hemisphere is equal to untreated controls **(Supplemental Fig. 1g)**. In contrast, total nodes in the ipsilateral hemisphere are significantly reduced relative to untreated controls after injection of either AAV8-*hSyn-mCherry* or AAV1-*pCAG-FLEX-EGFP* **(Fig. 1f)**. Node loss in both cases is evenly distributed between the apical and basal compartments **(Fig. 1g-h)**. To further understand how AAV-mediated gene delivery affects dendrites, we performed Strahler Order analysis ^40^. The maximum Strahler Order reached by a cell, termed its Strahler Number (SN), is a measure of branching depth, where higher numbers represent more extensively branched arbors **(Supplemental Fig. 2a)**. In untreated controls, 29% of neurons reach SN4, and similarly, 24%, or 32% of cells in the contralateral hemispheres reach SN4 after injection of AAV8-*hSyn-mCherry* or AAV1-*pCAG-FLEX-EGFP*, respectively. In striking contrast, for cells from the injected hemisphere, only 4% (after AAV8-*hSyn-mCherry*), or 7% (after AAV1-*pCAG-FLEX-EGFP*) reach SN4 **(Supplemental Fig. 2b-d)**. These results together reveal a dramatic loss of dendritic segments that underlies reduced dendritic length and complexity after administration of AAV.

High doses of AAV, which are often needed to drive robust transgene expression, can be neurotoxic ^41^. The dose of AAV8-*hSyn-mCherry* used here, 4×10^10^ vg, is near the middle of the range of viral titers commonly injected into the brain (~3.3×10^7^ - 2×10^13^ vg; reviewed in ^42^). To determine if reducing the number of viral genomes can prevent the effects of AAV8-*hSyn-mCherry* on dendrites, we injected a 1:20 dilution (2×10^9^ vg) of the same virus. With this promoter and system, 2×10^9^ vg is not sufficient to drive detectable *mCherry* expression **(Fig. 1a & Supplemental Fig. 1a)**. Dendritic length, Sholl intersections and total nodes are all significantly improved with this lower dose compared to a titer of 4×10^10^ vg and are indistinguishable from untreated controls **(Fig. 1i-l)**. However, only 14% of pyramidal cells in the ipsilateral hemisphere reach SN4, compared to 23% of cells in the non-injected contralateral hemisphere **(Supplemental Fig. 2e)**. Thus, a lower dose of AAV8-*hSyn-mCherry* can avoid some impacts on dendritic complexity, but is too low to drive detectable transgene expression. Overall, our results suggest that experimentally and clinically useful doses of AAV can substantially simplify and shrink dendrites.

### AAV1-*pCAG-FLEX-EGFP* disrupts synaptic transmission

The reduction in dendritic size and complexity after administering AAV1-*pCAG-FLEX-EGFP* **(Fig. 1)** significantly reduces the area over which these neurons can receive synaptic inputs. If AAV-induced dendritic loss is associated with a decrease in the number of functional synapses, it should be apparent as a reduction in the frequency of mEPSCs, which represent the postsynaptic response to spontaneous release of neurotransmitter at individual synapses. Consistent with loss of synapses, whole-cell patch clamp recordings reveal that mEPSC frequency is reduced, and inter-event interval is increased **(Fig. 2a-c)** in pyramidal cells from the AAV1-*pCAG-FLEX-EGFP*-injected hemisphere relative to untreated controls. mEPSC amplitude is also decreased in AAV-injected hemispheres **(Fig. 2d)** and rise and decay times are longer **(Fig. 2e-g)**. The slower kinetics of mEPSCs recorded in neurons from AAV-injected animals cannot be explained by their smaller peak amplitudes or smaller dendritic arbors, but could occur as the result of greater Ca^2+^ influx through AMPA receptors. AMPA receptors are heterotetramers, and receptors containing the GluA2 subunit are Ca^2+^ impermeable, while GluA2-lacking receptors allow Ca^2+^ influx. To determine the fraction of the whole cell current that is mediated by Ca^2+^-permeable AMPA receptors, we bath applied 1-Naphthylacetyl spermine (NASPM), a synthetic analog of Joro spider toxin that selectively blocks GluA2-lacking, Ca^2+^-permeable AMPA receptors in a use- and voltage-dependent manner ^43^. As expected, NASPM blocked only a small fraction (5.1%) of the spontaneous excitatory postsynaptic currents (sEPSC), recorded at −70 mV in untreated controls. In contrast, NASPM blocked 21.8% of the sEPSC in AAV-injected hemispheres, suggesting a greater than 4-fold increase in the fraction of the current carried by Ca^2+^-permeable AMPA receptors after AAV injection **(Fig. 2h-i)**. To independently assess the fraction of Ca^2+^-permeable AMPA receptors, we measured relative levels of GluA1 vs. GluA2 subunits in cortical lysates, since Ca^2+-^permeable AMPA receptors lack the GluA2 subunit. Following injection of AAV1-*pCAG-FLEX-EGFP*, levels of GluA2 are reduced relative to GluA1, consistent with a shift towards more Ca^2+^-permeable AMPA receptors **(Fig. 2j)**. Taken together, these data suggest that AAV-mediated gene delivery may cause a drop in synapse density that parallels the loss of dendritic length and disrupts synaptic transmission at remaining synapses.

**Figure 2.**
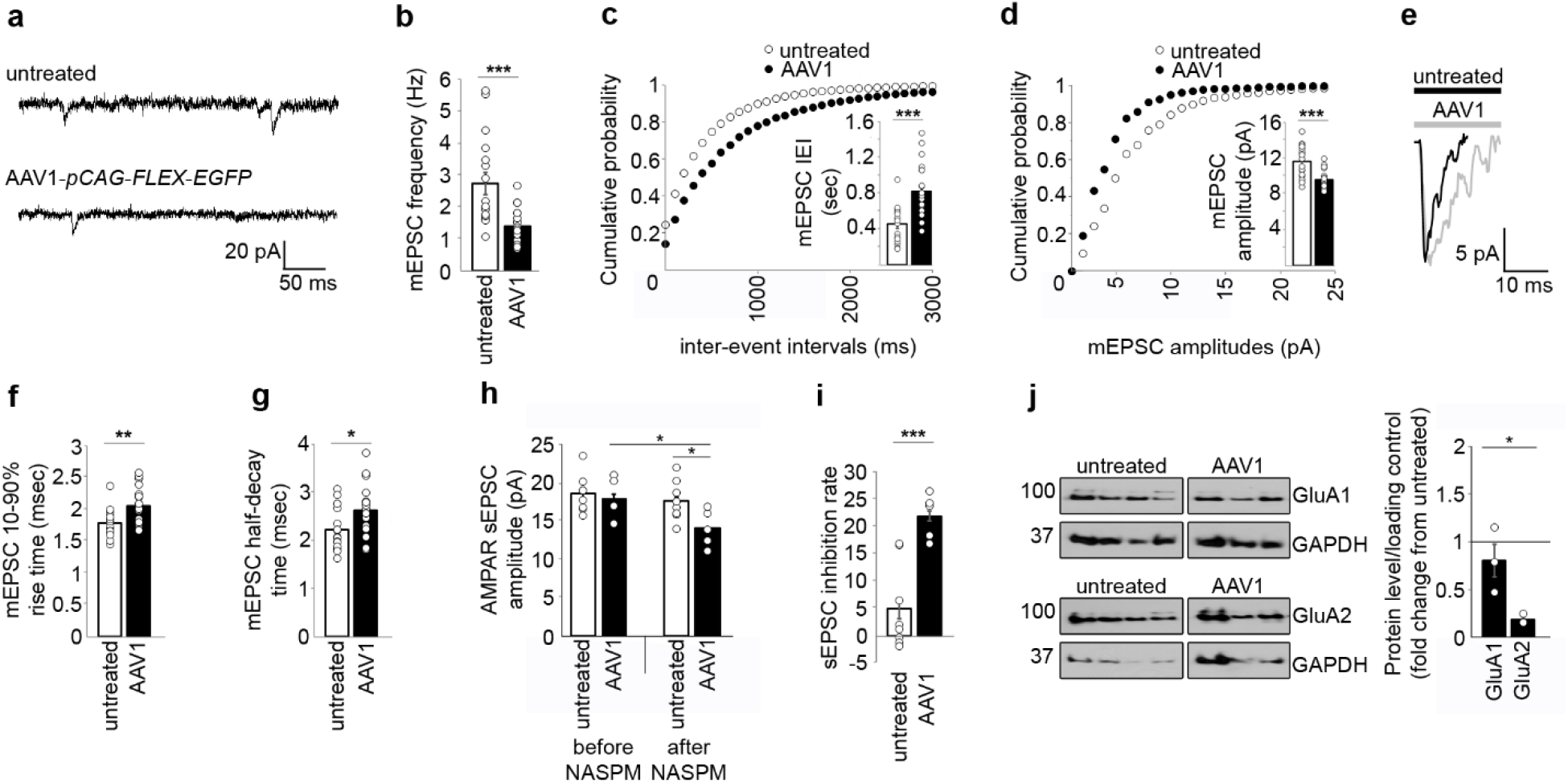
AAV1-*pCAG-FLEX-EGFP* in the absence of CRE disrupts excitatory synaptic transmission in somatosensory cortex three weeks post-injection. **(a)** Example whole cell recordings from S1 pyramidal cells in untreated controls or the ipsilateral hemisphere after AAV1 injection (4.2×10^9^ vg, as in Figure 1). **(b-c)** AAV1 administration reduces mEPSC frequency and increases the inter-event interval. Inset in **c**, mean and individual IEIs. **(d)** AAV1 is associated with a decrease in mEPSC amplitude. Inset, mean and individual amplitudes. **(e)** Representative mEPSCs from untreated controls (black) and cells ipsilateral to AAV1 injection (grey). **(f-g)** AAV1 increases mEPSC rise time and half-decay time. **(h-i)** As expected, NASPM does not significantly change sEPSC amplitude in untreated controls. However, the same treatment significantly decreases sEPSC amplitude in AAV1 injected samples. **(j)** AAV1 selectively decreases GluA2 levels, reducing the ratio of GluA2 to GluA1. **(a-g)** *n*=17 cells for untreated control; *n*=20 cells for AAV1. **(h-i)** *n*=8 cells for untreated control; *n*=5 cells for AAV1. **(i)** *n*=4 animals for untreated control; *n*=3 animals for AAV. Bars, mean ± S.E.M. *p<0.05; **p<0.01; ***p<0.001.

### AAV triggers persistent increases in immune gene expression

Because several immune proteins can regulate neuronal morphology and synaptic transmission ^29–37,44^, we wondered if AAV-associated dendritic loss could be a result of the host immune response. To explore this possibility, we examined the expression of a small subset of immune genes that can also regulate dendritic morphology (*C3, H2-K, H2-D*, and *Fyn*). To determine if any of these pleiotropic immune molecules are rapidly upregulated in response to AAV, we measured the levels of their mRNAs in samples of somatosensory cortex 4 days post-injection via RT-qPCR. To verify successful viral transduction at this time point, we injected AAV8-*hSyn-mCherry* and detected *mCherry* mRNA **(Supplemental Fig. 3a)**. In extracts of AAV8-*hSyn-mCherry* injected hemispheres, mRNAs encoding *H2-K, H2-D*, and *C3* are all significantly elevated relative to untreated controls, revealing an early transcriptional response to AAV-mediated gene delivery. Like dendritic loss, increases in immune gene mRNA levels are restricted to the injected hemisphere **(Supplemental Fig. 3b-e)**. Thus, AAV-mediated gene delivery rapidly upregulates the levels of immune proteins that can regulate dendritic complexity and synaptic transmission.

Dendritic loss and synaptic changes are apparent three weeks post-injection of AAV. To determine if AAV persistently upregulates levels of these same immune proteins, we performed Western blotting on lysates of injected somatosensory cortex three weeks post-injection. Sham surgery has no effect on the levels of these immune proteins and vehicle injection reduces H2-K levels **(Fig. 3b-d)**. In contrast, both AAV8-*hSyn-mCherry* and AAV1-*pCAG*-FLEX-*EGFP* significantly upregulate H2-K and H2-D relative to untreated and vehicle controls **(Fig. 3b-c)**. Fyn is persistently upregulated by AAV8-*hSyn-mCherry* but not AAV1-*pCAG-FLEX-EGFP*, and C3 is not increased by either virus at this time point. Persistent upregulation of immune proteins following AAV injection is not a unique feature of somatosensory cortex, because injection of AAV8-*hSyn-mCherry* into motor cortex and cerebellum locally upregulates H2-K, C3, and Fyn protein levels in both areas, relative to untreated and vehicle controls three weeks post injection **(Supplemental Fig. 4b-c, f-g)**. In somatosensory cortex, levels of H2-D (and to a lesser extent, H2-K) proteins inversely correlate with dendritic length across four conditions (untreated, vehicle, AAV8-*hSyn-mCherry*, or AAV1-*pCAG-FLEX-EGFP*) three weeks after injection, while C3 and Fyn levels do not correlate with dendritic length at this time point **(Fig. 3e)**. We observed a relatively small amount of mCherry fluorescence in fibers outside the injection areas, including in caudal regions of contralateral S1 and ipsilateral hippocampus within the stratum lacunosum-moleculare (**Supplemental Fig. 5a, d**). However, the presence of mCherry was not associated with changes in the levels of H2-K, H2-D, Fyn, or C3 in either region (**Supplemental Fig. 5b-c, e-f**). Together, these results show that dendritic loss induced by AAV-mediated gene delivery is associated with rapid and persistent upregulation of multiple immune proteins that can negatively regulate neurite length.

**Figure 3.**
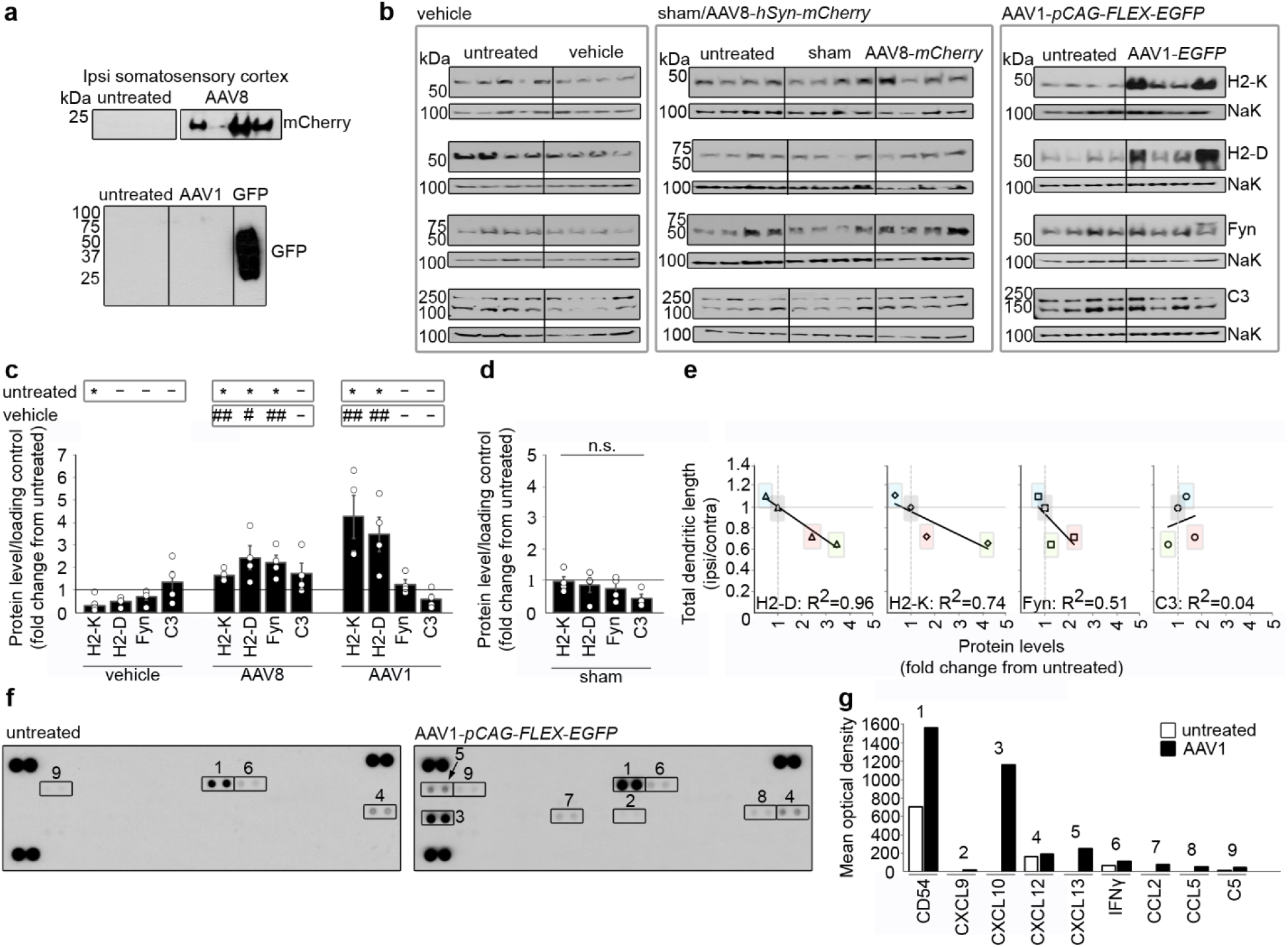
Administration of AAV causes persistent upregulation of diverse inflammatory mediators in somatosensory cortex three weeks post-injection. **(a)** mCherry positive immunoblot demonstrates successful viral transduction and transgene expression for AAV8. AAV1 (in the absence of CRE) shows no detectable EGFP expression in immunoblots. Positive control, pure GFP-GCaMP fusion protein. **(b)** Immunoblots used for analysis in **c-d**. **(c)** AAV8 (1.2×10^11^ vg) upregulates MHCI (H2-K, H2-D) and Fyn relative to untreated controls and vehicle. AAV1 (4.2×10^9^ vg) in the absence of CRE specifically upregulates H2-K and H2-D. **(d)** Sham surgery does not alter immune protein levels. **(e)** H2-D (triangle) and H2-K (diamond) levels, but not Fyn (square) or C3 (circle) levels, negatively correlate with dendritic complexity across treatments. Levels of different proteins for single treatment conditions (untreated (grey), vehicle (blue), AAV8 (red), or AAV1 (green)) are enclosed in shaded rectangles. **(f-g)** Cytokine array panels show upregulation of CD54, CXCL9, CXCL10, CXCL13, IFNγ, CCL2, CCL5 and C5 in ipsilateral somatosensory cortex 3 weeks after injection of AAV1-*pCAG-FLEX-EGFP* (4.2×10^9^ vg). *n*=4 mice per condition. Bars in **c** and **d** are mean ± S.E.M, rooted in untreated control (*p<0.05 relative to untreated control; #p<0.05; ##p<0.01 relative to vehicle). Loading controls in **b** (bands below each immune protein), Na^+^K^+^ ATPase.

In the current study, dendritic loss and upregulation of immune molecules are seen with viruses that have different capsid serotypes, encoded transgenes, and promoters **(Fig. 1)**, raising the possibility that these phenotypes are responses to a common feature of divergent AAVs. One feature that is shared by nearly all AAVs is a DNA genome that is rich in unmethylated CpG motifs, which are a feature of microbial DNA that is detected by the innate immunoreceptor TLR9. TLR9 activation leads to the release of proinflammatory cytokines that in turn upregulate MHCI and components of the complement cascade, including C3, paralleling what we see after AAV transduction ^6–10,26,27,45^. To determine if cytokine signaling downstream of TLR9 is elevated in the CNS following AAV injection, we used an unbiased ELISA-based array to simultaneously moniter the levels of forty different cytokines, chemokines, and acute phase proteins. Three weeks after injection of AAV1-*pCAG-FLEX-EGFP*, multiple cytokines and chemokines are upregulated relative to untreated controls, including IFNγ, CD54 (sICAM-1), complement component C5, CXCL9 (MIG), CXCL10 (IP-10), CXCL13 (BLC), CCL2 (JE) and CCL5 (RANTES) **(Fig. 3f-g)**. Notably, all of the upregulated cytokines and chemokines can be upregulated by TLR9 activation using CpG-rich DNA ^26,46,47^, and a subset (CD54 (sICAM-1), CXCL10 (IP-10), CCL2 (JE) and CCL5 (RANTES)) are upregulated in astrocytes and microglia after TLR9 activation ^47^. Together, our cytokine arrays, RT-qPCR, and Western blots demonstrate that AAV administration in the CNS causes increases in multiple innate and adaptive immune molecules that limit dendritic complexity and synaptic transmission. These results raise the possibility that some or all of these proteins could contribute to the morphological and/or functional changes we observe after AAV administration. A unifying feature of these many candidate mediators is that they can all be upregulated by activation of TLR9.

### Dendritic simplification is rescued by limiting detection of the AAV genome via TLR9

We find that AAV-mediated gene delivery is associated with significant loss of dendritic complexity and changes to the functional properties of remaining synapses. We also see the upregulation of many antiviral immune proteins that work in concert to maintain proper neuronal structure and function in the healthy nervous system, raising the possibility that the neuronal phenotypes we observe are the cumulative effect of disruptions to multiple neuronal homeostatic pathways. If this is the case, then targeting individual upregulated immune proteins is unlikely to fully prevent AAV-induced dendritic changes. Because the upregulated proteins can all be induced by TLR9 activation, we wondered if a TLR9 antagonist might be the most direct, effective, and simplest approach to prevent AAV-induced dendritic loss.

To limit initial detection of AAV’s unmethylated CpG-rich genome via TLR9, we administered a commercially available oligonucleotide antagonist of TLR9, CpG ODN 2088, one hour before and 24 hours after injection of 4.2×10^9^ vg of AAV1-*pCAG-FLEX-EGFP*. Systemic administration of an inactive analogue of ODN 2088 (ODN CTL) does not alter dendritic length or complexity in cortical pyramidal cells of S1 (not shown), indicating that injection of oligonucleotides is not sufficient to disrupt dendritic arbors. In addition, injection of AAV1-*pCAG-FLEX-EGFP* in the presence of ODN CTL reduced dendritic length and complexity in a manner that closely resembled changes caused by AAV1-*pCAG-FLEX-EGFP* alone **(Fig. 4a-c)**. In striking contrast to the loss of dendritic length and complexity caused by AAV1-*pCAG-FLEX-EGFP* alone or in the presence of ODN CTL, the TLR9 inhibitor ODN 2088 significantly improves total dendritic length and Sholl intersections to values that are indistinguishable from untreated controls **(Fig. 4a-c)**. CpG ODN 2088 also preserves Sholl intersections throughout the length of the dendritic arbor **(Fig. 4d)**. These results suggest that AAV’s genome is detected by TLR9 in the CNS and demonstrate that AAV-induced dendritic loss can be prevented using a TLR9 inhibitor.

**Figure 4.**
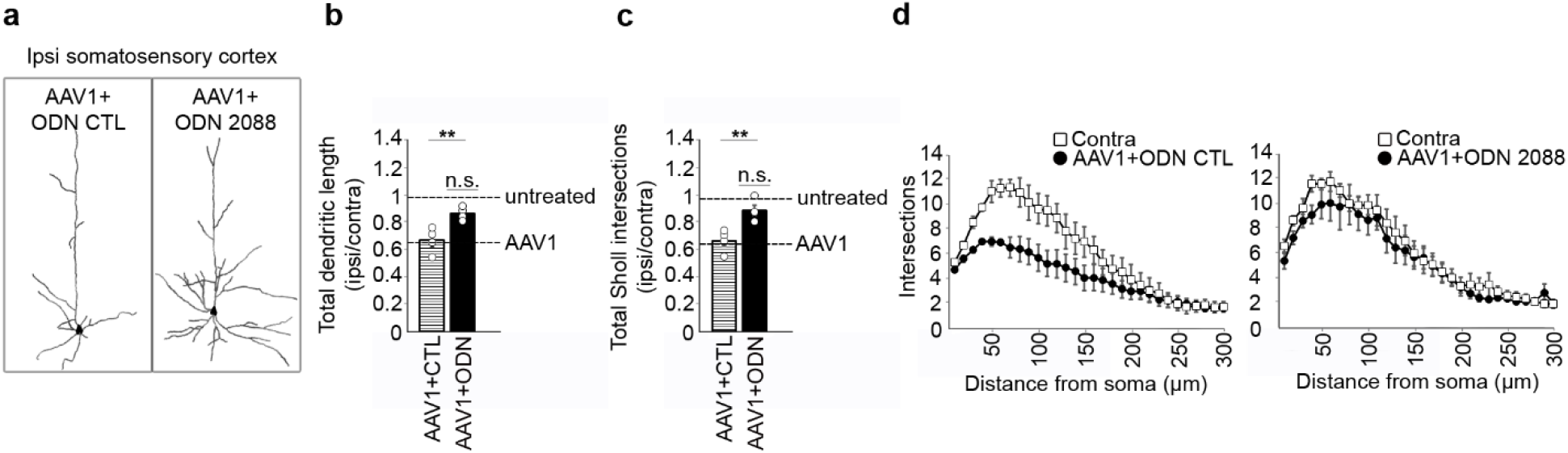
Loss of dendritic complexity can be prevented by limiting AAV detection using the TLR9 antagonist, CpG ODN 2088. **(a)** Example tracings of cells taken from the ipsilateral hemisphere of S1 after AAV1+ODN CTL or AAV1+ODN 2088 injection (4.2×10^9^ vg, as in Figures 1–3). **(b-c)** Total dendritic length and total Sholl intersections for AAV1+ODN 2088 are significantly higher than for AAV1+ODN CTL, and indistinguishable from untreated controls. **(d)** Protection in the presence of ODN 2088 is effective throughout the dendritic arbor. *n*=4 animals per treatment. Bars, mean ± S.E.M. **p<0.01 relative to AAV1+ODN 2088 CTL.

## Discussion

Here, we show that AAV-mediated gene delivery significantly reduces dendritic complexity and disrupts synaptic transmission in adult rodent cortex. Dendritic loss in somatosensory cortex is apparent three weeks after AAV injection and is accompanied by a decrease in the frequency and amplitude of mEPSCs and an increase in the proportion of Ca^2+^-permeable AMPA receptors. Dendritic simplification does not require transgene expression, is dose-dependent, and is not restricted to a specific AAV capsid serotype, promoter, preparation method, or encoded transgene. Disrupted neural homeostasis in AAV-injected animals is associated with the rapid and persistent upregulation of several immune molecules that can limit dendritic complexity, all of which can be upregulated by TLR9 activation, and blocking immunodetection of the AAV genome via TLR9 preserves dendritic arbors in AAV-injected animals. These results reveal previously unsuspected impacts of the immune response to AAV’s genome on circuit structure and function and identify practical approaches to improve the safety and efficacy of viral gene delivery to the nervous system.

The doses of AAV used here (2×10^9^ - 2.1×10^11^ vg) fall within the range of titers commonly used for gene delivery to the CNS in preclinical studies (~3.3×10^7^ - 2×10^13^ vg). In the clinic, doses are generally at or above the high end of this range (~1.2×10^10^ - 3.3×10^14^ vg; reviewed in ^42^). We find reducing the dose of AAV can avoid most elements of dendritic simplification. However, such a lower titer is insufficient to drive detectable *mCherry* expression in the current system. It is possible that titer windows can be found, particularly with strong promoters, that are low enough to avoid dendritic impacts, but still sufficient to drive functionally relevant levels of transgene expression. However, the appropriate multiplicity of infection will have to be individually determined for each unique combination of capsid serotype, promoter, encoded transgene, cell population and route of administration. At present, most experiments and clinical applications use relatively high levels of virus to ensure robust transgene expression, levels that are comparable to or above levels that give rise to severe dendritic deficits.

A major finding of this study is that in response to AAV, dendritic arbors of cortical pyramidal neurons become less complex, losing entire segments. Dendritic morphology powerfully influences integration and other emergent properties that arise due to the electrical separation of synapses, suggesting that AAV-driven dendritic loss is likely to have a potent effect on the computations and outputs of cortical pyramidal neurons. In addition to morphological changes, whole-cell patch clamp recordings reveal altered synaptic transmission in pyramidal neurons within AAV-injected hemispheres. The drop in dendritic size after AAV is accompanied by a drop in mEPSC frequency, consistent with the loss of functional synapses. Supporting the idea that the remaining synapses are functionally altered after AAV injection, there is an increase in the fraction of the sEPSC that is mediated by Ca^2+^-permeable AMPA receptors, and a concomitant decrease in expression of the AMPA receptor subunit GluA2, which confers Ca^2+^-impermeability. The combination of broad dendritic loss and altered AMPA receptor subunit composition together represents significant functional changes in cortical circuits following administration of AAV.

Many studies examining the safety of AAV-mediated gene delivery in the CNS have focused on classic hallmarks of inflammation, including the presence of infiltrating immune cells and neutralizing antibodies, viral clearance, and ultimately cell death. Our studies reveal that neuroimmune crosstalk mediates additional, unexpected effects of AAV on dendritic morphology. Because it is triggered by the detection of CpG-rich DNA, dendritic loss may occur wherever AAV genomes are present, even if transgene expression is far more restricted, e.g., due to CRE dependence or use of a cell-type specific promoter. These AAV-induced changes to dendritic structure and function may have gone undetected in studies that used empty or fluorophore-encoding vectors as negative controls, since AAV-induced TLR9 activation was present in both experimental and control conditions. Comparing empty vector to untreated or vehicle controls is one way to isolate and monitor the effects of AAV itself.

The experiments performed here indicate that AAV-induced TLR9 activation drives persistent dendritic loss following AAV transduction *in vivo*. The current findings complement and extend findings that other TLR9 agonists, including synthetic CpG-DNA, can cause neurite retraction *in vitro* ^48,49^. We also see upregulation of a wide range of innate and adaptive immune proteins that act downstream of TLR9 and can regulate dendritic complexity. Four days after injection of AAV8-*hSyn-mCherry*, mRNAs encoding *H2-K, H2-D*, and *C3* are upregulated in somatosensory cortex. These findings from mouse cortex align well with previous studies showing upregulation of MHCI four days after striatal injection of AAV9 or AAV2 in rats ^50,51^, and AAV-induced upregulation of *H2-D* in cortical microglia of mice three days post-injection ^52^. In addition to the rapid response, we see persistent upregulation of H2-K and H2-D proteins three weeks after injection of either AAV8-*hSyn-mCherry* or AAV1-*pCAG-FLEX-EGFP* in somatosensory cortex, and in AAV8-*hSyn-mCherry*-injected motor cortex and cerebellum. Finally, we see persistent upregulation of cytokines and chemokines that act immediately downstream of TLR9, including CD54, CCL2, CCL5, IFNγ, CXCL13, and CXCL10, in injected S1. Many of the upregulated immune proteins have additional roles in the CNS, where they negatively regulate neurite length, dendritic complexity and synaptic transmission ^28–38,44,53–60^. Future studies are needed to individually define the roles of each of the many upregulated immune proteins in AAV-induced disruptions to dendritic and synaptic function, to elucidate the role of non-neuronal cell types, including microglia and astrocytes, and to determine if similar changes are triggered by bacteria and other microbes that contain unmethylated CpG-rich DNA genomes. Because signaling by different members of the TLR family converge at key points, it will be of interest to determine if the RNA genomes of other commonly used viral vectors, such as lentivirus and pseudorabies virus, which are detected by TLR7, also induce immune responses that disrupt dendritic arbors. Overall, our studies show that upregulation of immune gene expression after AAV transduction is associated with structural and functional changes in pyramidal neurons, and that immunodetection of AAV’s genome by TLR9 is a central event that triggers AAV-induced dendritic loss.

Recombinant AAVs are powerful tools for gene delivery and have emerged as the vector of choice in the nervous system. Despite an overall strong safety profile, AAV-mediated gene delivery has been associated with neurotoxicity in specific cell populations, some of which can be attributed to transgene expression. AAV-mediated gene delivery into the CNS of pigs or non-human primates causes dorsal root ganglion (DRG) toxicity that is alleviated by knocking down transgene expression ^61,62^. Additionally, overexpression of SMN1 in a spinal muscular atrophy model caused a toxic gain-of-function in mice ^63^. These and other studies suggest that the levels and distribution of transgene products can be relevant to avoiding neurotoxicity after AAV-mediated gene delivery. However, several lines of evidence suggest that the changes in dendritic structure seen here cannot be ascribed to overexpression of a foreign or toxic transgene. First, AAV1-*pCAG-FLEX-EGFP* causes severe deficits in the absence of any detectable transgene expression, even after antibody amplification. Second, in our analysis of AAV8-*hSyn-mCherry* injected animals, we generally did not trace cells that expressed readily detectable *mCherry*, but still observed significant simplification relative to cells of the contralateral hemispheres. Third, dendritic complexity in contralateral S1 is indistinguishable from untreated controls, despite the presence of mCherry that is trafficked from the injected hemisphere. Together these findings demonstrate that transgene expression is neither necessary nor sufficient for AAV-associated dendritic loss. In contrast, our studies with ODN 2088 demonstrate that TLR9 activation is required for dendritic loss following injection of AAV, supporting a central role for immunodetection of the viral genome in dendritic disruption.

Like transgene expression, the presence of AAV genomes has been linked to neuronal cell death. Broadly active *cis*-regulatory elements are associated with dose-dependent, cell-type specific toxicity after subretinal injection of AAV8, AAV5, or an AAV2 in mice, and toxicity is still observed with AAVs that encode no transgene ^17^. Furthermore, AAV-associated retinal toxicity can be either prevented or delayed by inhibiting detection of the genome via TLR9 in multiple mammalian model systems ^19^. Electroporating only AAV’s ssDNA inverted terminal repeats (ITRs) into progenitor cells *in vitro* recapitulates dose-dependent loss of neural progenitor cells and immature adult-born dentate granule cells following injection of AAV into dentate gyrus ^18^. This ITR-driven toxicity does not require Sting, a TLR9-independent pathway for detecting foreign DNA. It will be of interest to determine if AAV-associated cell death in the dentate gyrus, like AAV-associated cell death in retina ^19^, and the AAV-induced dendritic loss we see in cortex, can be improved by inhibition of TLR9.

We find that several different approaches may allow researchers to minimize or avoid AAV-induced dendritic simplification. First, reducing the number of viral genomes injected can reduce AAV-induced loss of dendritic complexity. However, this method also reduces transgene expression, which may not be desirable in experimental or clinical settings. A second approach is to inject AAV into areas that are directly connected to the target of interest and allow the transgene to travel intracellularly to achieve local expression. This approach is facilitated by the creation of AAVs with specific transport properties, such as AAV-retro. A recent study that identified AAV-induced cell death in dentate gyrus was able to achieve safe hippocampal transgene expression by injecting an AAV-retro serotype into downstream targets, and allowing the virus to backpropagate through axonal projections ^18^. We see transgene expression in fibers in contralateral S1, which does not show upregulation of immune genes or changes in dendritic structure, suggesting that remote injection can protect dendrites while permitting local transgene expression in projection fibers. Finally, we show that AAV-induced dendritic simplification can be avoided using systemic injection of a TLR9 antagonist, ODN 2088. Because ODN 2088 is a short oligonucleotide, it can be introduced into host cells as part of the viral genome, *in cis*, an approach that has recently been used to reduce or delay AAV-induced toxicity in the retina in pigs and macaques, respectively ^19^. A significant advantage of using TLR9 inhibitors is that it does not require a reduction in effective viral dosage, either through dilution of injected virus or indirect injection of distant sites. Together, these studies reveal unexpected effects of AAV-mediated gene delivery on neuronal structure and function and identify the immune response to the AAV genome as the triggering event. These studies also identify TLR9 inhibitors as an important strategy to avoid AAV-induced dendritic loss and improve the safety and efficacy of AAV-mediated gene therapy in the nervous system.

## Acknowledgements

We wish to thank Esteban Engel and Elise Cope for their technical assistance during these experiments.

## Author Contributions

C.S. and L.B. conceived the study and wrote the paper. C.S. designed figures and performed data analysis, western blots, fluorescent imaging and DiO labeling. N.K. performed blinded dendritic Sholl and Strahler Order analysis. J.M. performed electrophysiology experiments. J.V. and C.J. performed AAV injections and collected tissue for analysis.

The authors declare no competing interests with the production of this article.

## Tables

**Supplemental Table 1.**
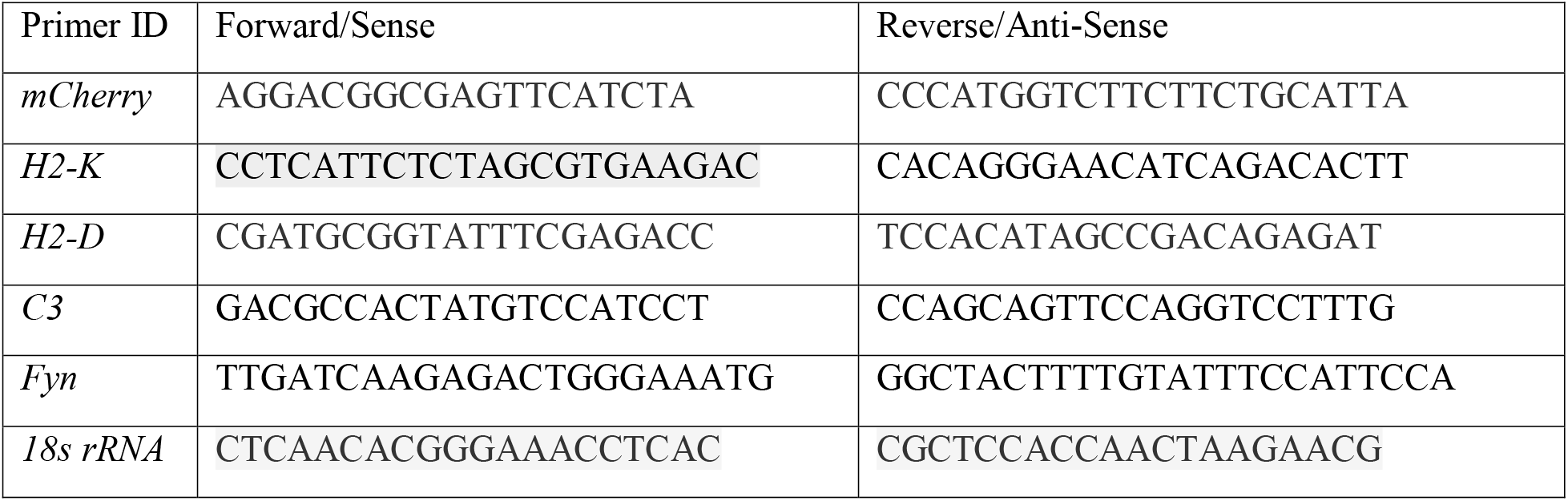
Primers used for RT-qPCR, four days post-injection.

**Supplemental Table 2.** Antibodies used for Western blotting, three weeks post-injection.

| Antibody | Supplier | Dilution |
| --- | --- | --- |
| Mouse $\alpha$ Na<sup>+</sup>K<sup>+</sup> ATPase | *Santa Cruz Biotechnology Catalog # sc-71638* | 1:50,000 |
| P8 (rabbit $\alpha$ H2-K) | *Genscript USA inc. (custom)* | 1:1,000 |
| Mouse $\alpha$ H2-D<sup>b</sup> | *Abcam Catalog # ab25228* | 1:250 |
| Mouse $\alpha$ Fyn | *Santa Cruz Biotechnology Catalog # sc-434* | 1:500 |
| Rabbit $\alpha$ C3 | *Abcam Catalog # ab200999* | 1:2,000 |
| Rabbit $\alpha$ RFP | *Rockland Antibodies and Assays Catalog # 600-401-379* | 1:500 |
| Rabbit $\alpha$ GFP | *Abcam Catalog # ab290* | 1:10,000 |
| Mouse $\alpha$ GFP | *Novus Biologicals Catalog # NBP2-22111SS* | 1:4,000 |
| Goat $\alpha$ Mouse IgG | *Invitrogen Catalog # A24512* | 1:5,000 |
| Goat $\alpha$ Rabbit IgG | *Jackson Laboratories Catalog # 111-035-046* | 1:5,000 |

---

**Supplemental Figure 1.**
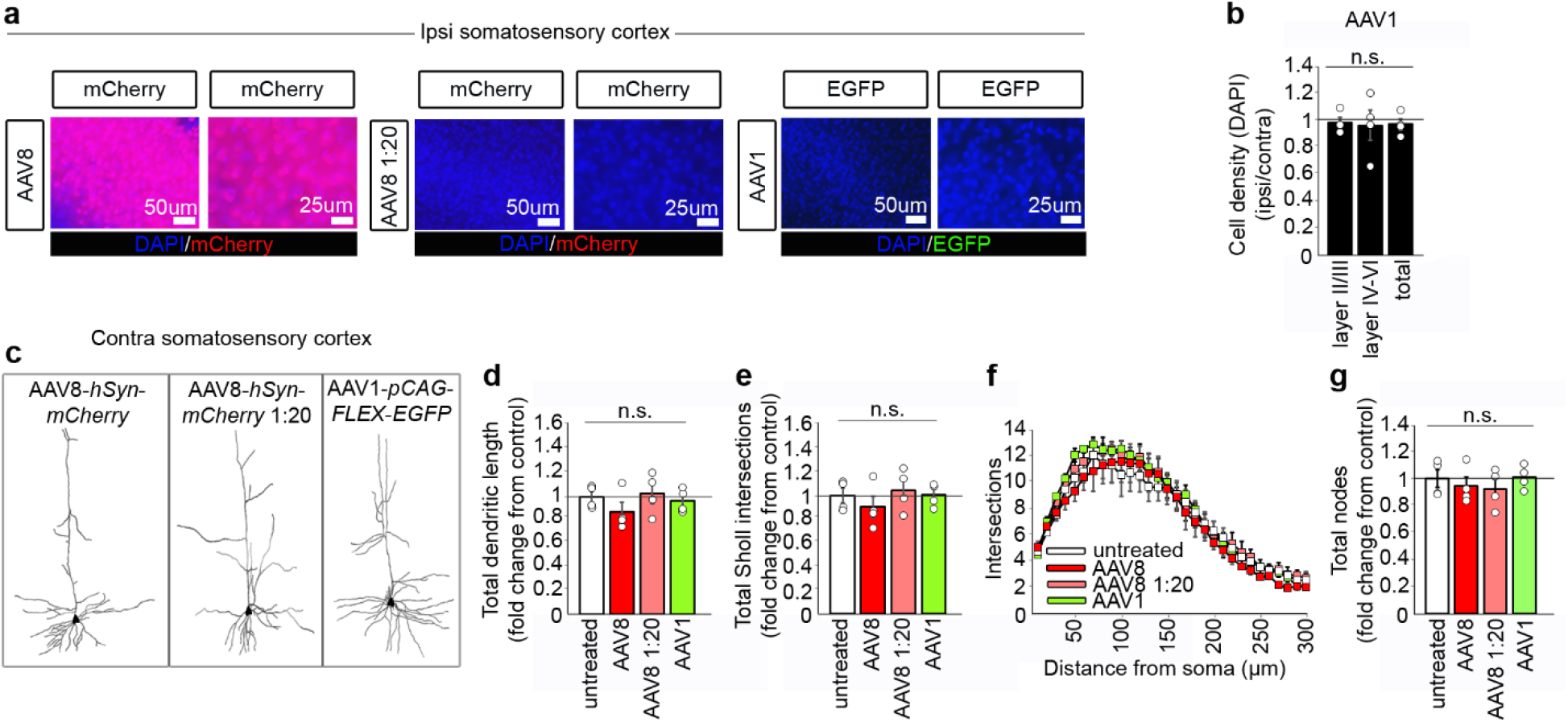
Dendritic complexity in the non-injected contralateral hemisphere is not affected by AAV-mediated gene delivery. **(a)** mCherry-positive cells in ipsilateral S1 after injection of AAV8-*hSyn-mCherry* (4×10^10^ vg; left), contrasting with lack of transgene expression after injection of a 1:20 dilution of the same virus (2×10^9^ vg; middle) or after injection of AAV1-*pCAG-FLEX-EGFP* (4.2×10^9^ vg; right). **(b)** Cell density counts from ipsilateral and contralateral hemispheres after injection of AAV1-*pCAG-FLEX-EGFP* (Princeton Viral Vector Core, Princeton NJ) in layers II/III, IV-VI and II-VI. **(c)** Example tracings of pyramidal cells from non-injected contralateral hemispheres. **(d-g)** Data from S1 pyramidal cells contralateral to AAV injection. Dendritic length, Sholl intersections and total nodes are indistinguishable from control. Bars, mean ± S.E.M, rooted in untreated control.

**Supplemental Figure 2.**
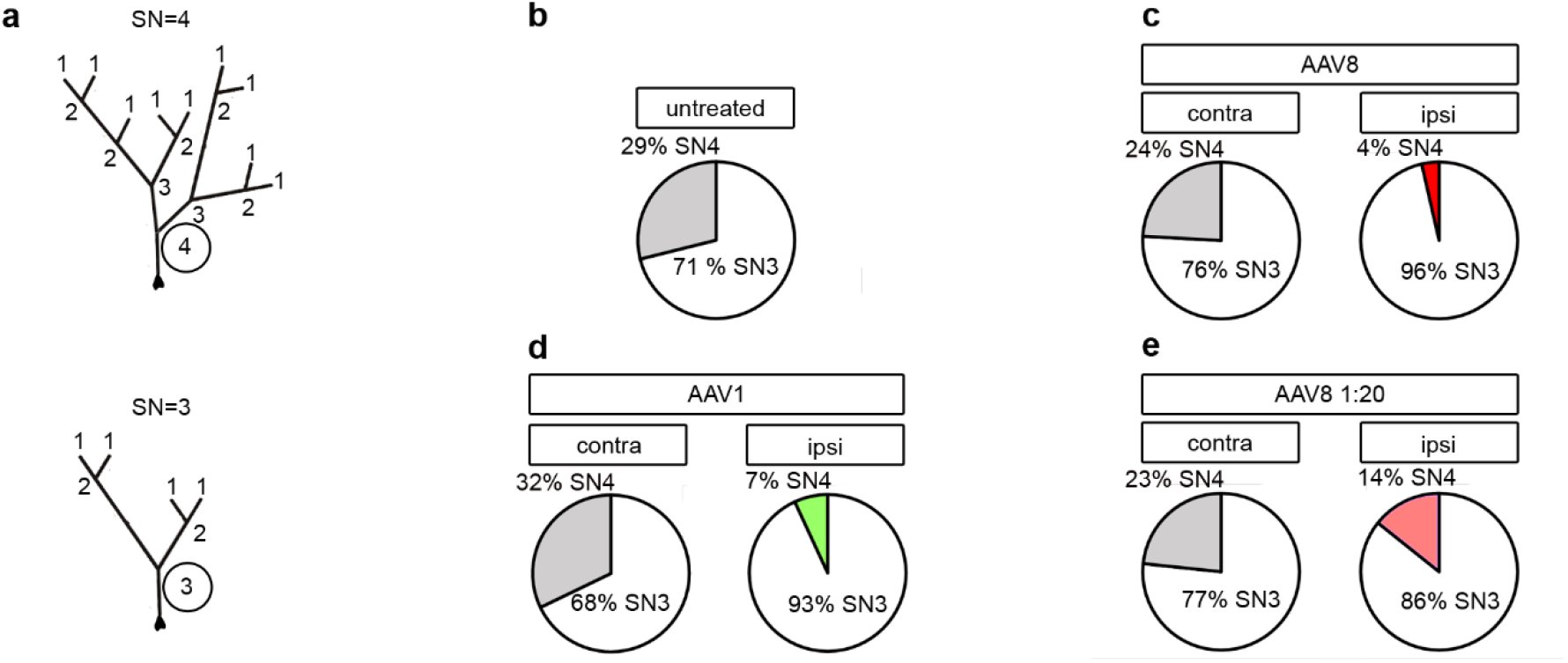
AAV reduces Strahler Number. **(a)** Example Strahler Order (SO) labeling of model dendrites with a Strahler Number (SN) of either 4 or 3. **(b-e)** Percentage of cells that reach SN 4 or 3 in untreated controls, or contralateral and ipsilateral hemispheres after AAV8 or AAV1 injection. Shaded or colored wedges represent more complex neurons with SN of 4. Total cells: untreated, 46; AAV8 contra, 29; AAV8 ipsi, 28; AAV8 1:20 contra, 30; AAV8 1:20 ipsi, 28; AAV1 contra, 28; AAV1 ipsi, 29.

**Supplemental Figure 3.**
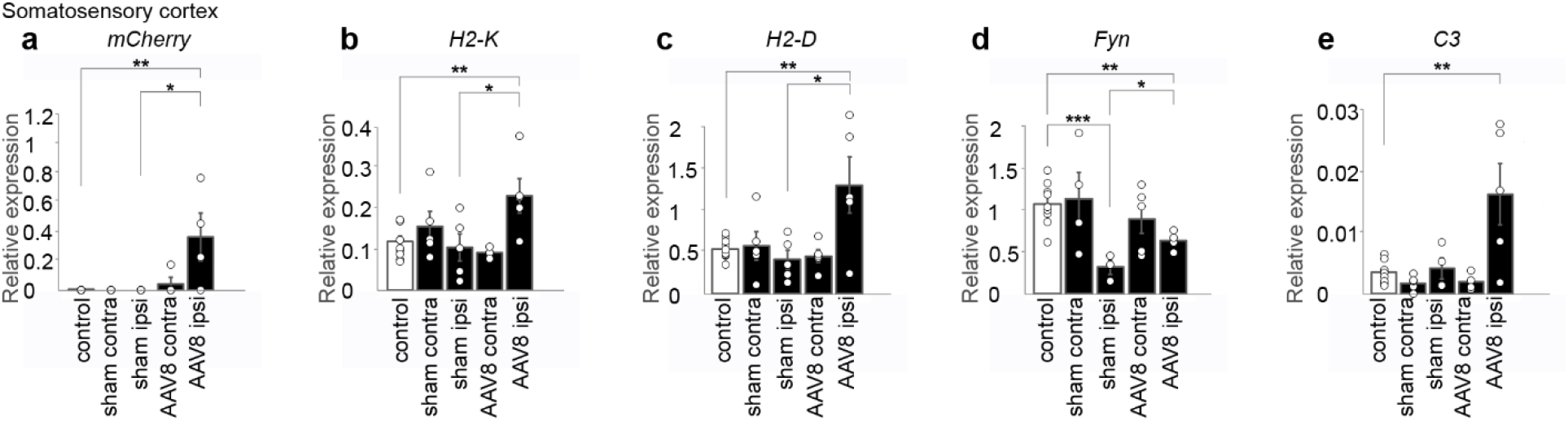
AAV8-*hSyn-mCherry* rapidly upregulates immune gene expression four days post-injection in somatosensory cortex. **(a)** *mCherry* mRNA detected in somatosensory cortex. (**be**) AAV8-*hSyn-mCherry* upregulates *H2-K, H2-D* and *C3* gene expression in injected hemisphere relative to untreated controls. *H2-K, H2-D* and *Fyn* levels are also upregulated after AAV8 injection relative to sham surgery. Injected, 1.26×10^11^ vg. Bars, mean ± S.E.M. (*p<0.05; **p<0.01; ***p<0.001). All values are normalized to a single reference gene (*18s rRNA*).

**Supplemental Figure 4.**
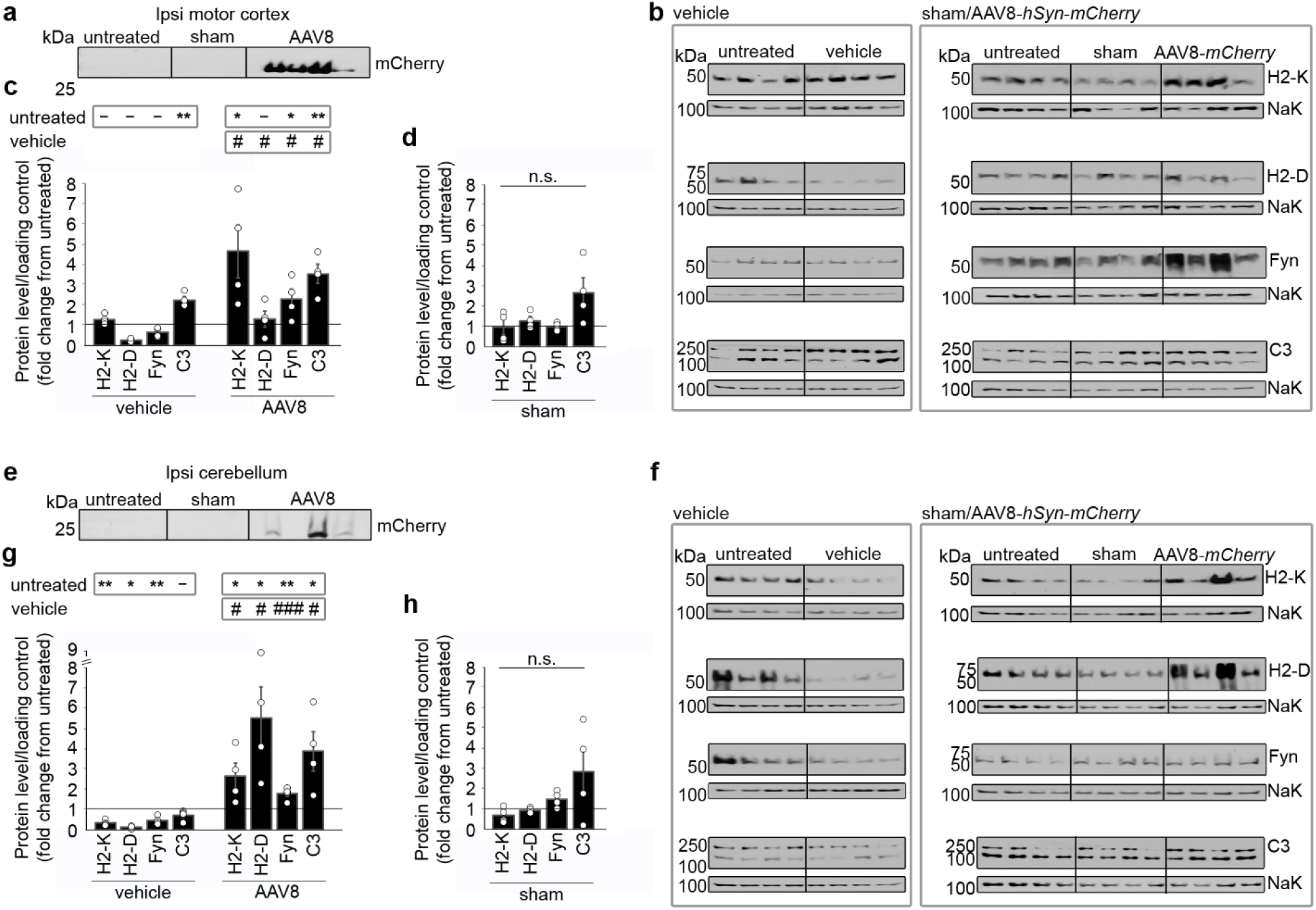
AAV-mediated gene delivery to motor cortex or cerebellum upregulates immune protein levels three weeks post-injection. **(a)** mCherry-positive immunoblot of motor cortex homogenates following injection of 1.26×10^11^ vg of AAV8-*hSyn-mCherry*, indicating successful viral transduction. **(b-c)** AAV8-*hSyn-mCherry* upregulates H2-K, Fyn and C3 levels relative to both vehicle and untreated controls. **(d)** Sham surgery does not alter protein levels in motor cortex. **(e)** mCherry positive immunoblot of cerebellum homogenates following injection of 1.26×10^11^ vg of AAV8-*hSyn-mCherry*, indicating successful viral transduction. **(f-g)** AAV8-*hSyn-mCherry* upregulates H2-K, H2-D, Fyn and C3 relative to both vehicle and untreated controls. **(h)** Sham surgery does not alter protein levels in cerebellum. *n*=4 mice for each condition. Bars, mean ± S.E.M rooted in untreated control (*p<0.05; **p<0.01 relative to untreated control; #p<0.05; ##p<0.01; ###p<0.001 relative to vehicle). Loading control, Na^+^K^+^ ATPase (below each immune protein).

**Supplemental Figure 5.**
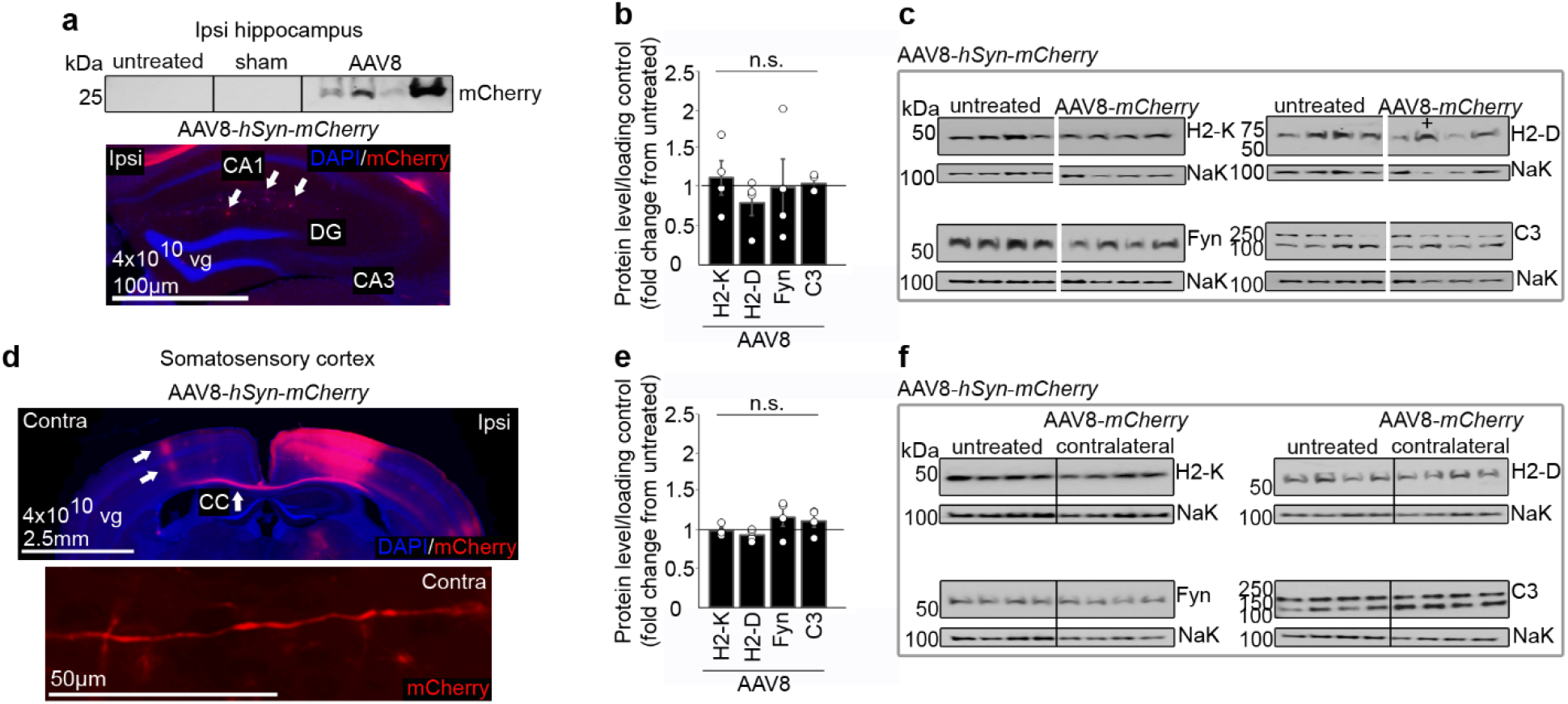
Despite presence of mCherry in non-injected areas of AAV-injected brain, immune proteins are not upregulated in these regions. **(a-c)** Hippocampus immediately ventral to injected cortex. Fluorescent mCherry protein is present in fibers in stratum lacunosum-moleculare of the hippocampus (white arrows in **a**) and is detectable in immunoblots of hippocampal lysates (**a**) but is not associated with altered immune protein levels (**b-c**). **(d-f)** mCherry is present in fibers in non-injected contralateral S1 (unlabeled white arrows in **d**), but immune protein levels are unaltered (**e-f**). **d** bottom panel, mCherry fluorescence in a fiber from area of the lower arrow in contralateral hemisphere. + in **c** represents one sample where a different exposure of the blot was used for analysis of both sample and loading control, to ensure values remained in the linear detection range. Bars, mean ± S.E.M rooted in untreated control. CA1, cornus ammonis 1; CA3, cornus ammonis 3; DG, dentate gyrus; CC, corpus callosum. Loading control, Na^+^K^+^ ATPase (NaK, bottom row of each blot).

**Supplemental Figure 6.**
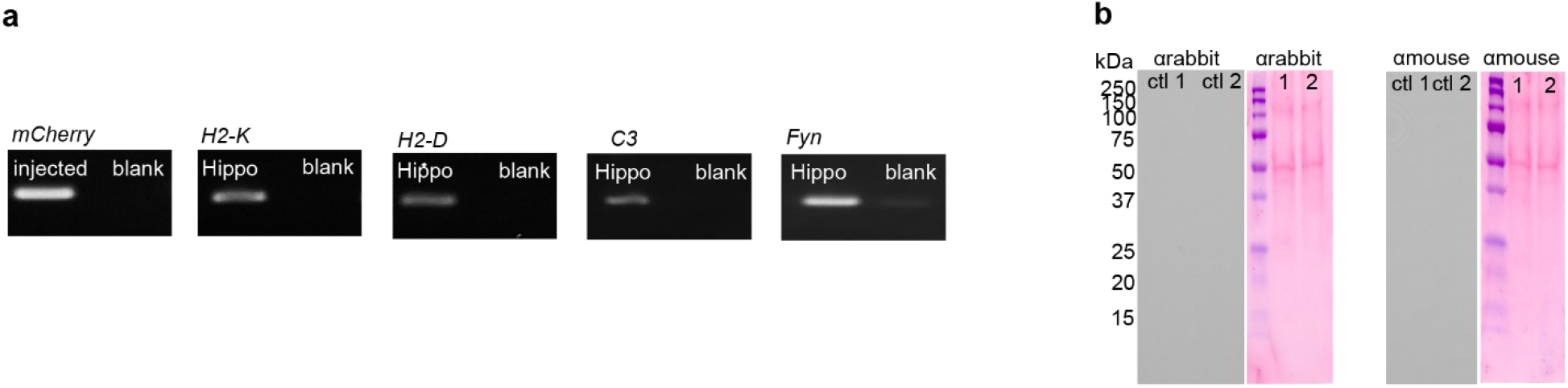
Negative controls for primers used in RT-qPCR, and antibodies used in immunoblots. **(a)** Blank lanes contained no sample, and lack of band in blank lane indicates absence of primer-dimers. **(b)** No primary negative controls for antibodies used in Figures 2–3 and Supplemental Figures 4–5. Ponceau stain of the same membranes shows protein was present in lanes used for immunoblots.

## Online Methods

### 2.1 Animals

Male C57BL/6J mice, 8 to 10 weeks old, were obtained from Jackson Laboratory (The Jackson Laboratory, Bar Harbor, ME). All mice acclimated at least 48 hours in the Princeton Neuroscience Institute vivarium before experiments were performed. All experiments were performed on animals 11 to 14 weeks old unless otherwise noted. Animals were group housed in Optimice cages (Animal Care Systems, Centennial, CO) with blended bedding (The Andersons, Maumee, OH) and enrichment (paper nesting and heat-dried virgin pulp cardboard hut). All mice were maintained on a 12hr light-dark cycle with ad libitum access to food (PicoLab Rodent Diet food pellets, LabDiet, St. Louis, MO) and water. All procedures were performed in accordance with protocols approved by the Princeton University Institutional Animal Care and Use Committee (IACUC) and in accordance with the animal welfare guidelines of the National Institutes of Health.

### 2.2 AAV Administration

Surgeries were performed in accordance to previously published procedures ^64^. Mice were anesthetized with isoflurane (5% for induction, followed by 1-2%, in oxygen; 1 L/min; Viking Medical) and mounted into a stereotaxic instrument (David Kopf Instruments, Tujunga, CA). Body temperature was maintained and monitored using Kent Scientific PhysioSuite (Kent Scientific Corporation, Torrington, CT). An osmotic diuretic drug (15% D-mannitol in DPBS, 0.25 ml) was administered via intraperitoneal injection ten minutes prior to craniotomy.

Viruses were prepared using endotoxin-free plasmid prep kits, and virus preps were endotoxin-negative. Unless otherwise noted, 4×10^10^ or 2×10^9^ total viral genomes (vg) of AAV2/8-*hSyn-mCherry-WPRE-hGH-pA* (Princeton Viral Vector Core, Princeton, NJ **(Fig. 1, 3; Supplemental Fig. 1–5)**, or 4.2×10^9^ vg of AAV2/1-*pCAG-FLEX-EGFP-WPRE* (UPenn; Addgene # 51502 **(Fig. 1–4; Supplemental Fig. 1–2)** or Princeton Viral Vector Core, Princeton, NJ **(Supplemental Fig. 1b)**) were unilaterally injected to brain regions using borosilicate glass capillaries pulled on a Sutter Micropipette Puller (Model P-2000, Sutter Instrument Company). Animals that received the AAV2/1-*pCAG-FLEX-EGFP-WPRE* virus did not receive any form of CRE recombinase, and therefore should not express EGFP. Lack of EGFP was confirmed by both confocal immunofluorescence microscopy **(Fig. 1a & Supplemental Fig. 1a)** and by using immunoblotting to amplify EGFP signals **(Fig. 3a)**. To ensure robust viral transduction, a total volume of 200 nanoliters (nl) was injected over 3 depths (500, 250, and 150 μm below the dura, 67 nanoliters per depth) in regions corresponding to the motor (+2.46 to +1.42 mm from Bregma) and somatosensory cortex (+0.2 to −0.8 mm from Bregma and 2 mm lateral) and cerebellum (−5.40 to −8.24 mm from Bregma) of the left hemisphere. For rescue experiments, 100 μg/25 g body weight of the Tolllike receptor 9 (TLR9) antagonist, CpG oligodeoxynucleotide (ODN) 2088, or of control oligonucleotide (ODN 2088 CTL), were injected intraperitoneally 1 hour before and 24 hours after AAV2/1-*pCAG-FLEX-EGFP* injection. All mice received a non-steroidal anti-inflammatory drug, Rimadyl (carprofen, Zoetis, Florham Park, NJ), immediately post-surgery and 24 h post-surgery (0.2 ml, 50 mg/ml, subcutaneous). Vehicle and sham surgery animals underwent the same procedures. Vehicle injected animals received 200 nl of buffer used to deliver AAV (PBS, 35 mM sodium chloride, 0.001% pluronic F-68 and 5% glycerol) and sham surgery mice did not receive any fluid injection. Untreated controls received no surgery or drug administration.

### 2.3 Diolystic Labeling and Sholl Analysis

Diolystic labeling and Sholl analyses to quantify pyramidal cell morphology were performed three weeks after AAV injection. Mice were anesthetized and intracardially perfused with 1.5% PFA diluted in PBS (Millipore Sigma Catalog # P3813-10PAK), and whole brains were removed. Brains were then post-fixed for one hour in 1.5% PFA and stored in cold PBS until used (usually 1-2 weeks). Brains were sectioned by vibratome (Leica Biosystems) at 100 μm and sections were stored in cold PBS. Brains that were injected with vehicle, AAV1-*pCAG-FLEX-EGFP*, or the 1:20 dilution of AAV8-*hSyn-mCherry* were notched with a scalpel after fixation on the lower half of the injected hemisphere, to allow identification of the injected hemisphere in the absence of transgene expression. Notching fixed sections does not alter dendritic complexity, because both vehicle control and AAV8 1:20 titer cohorts were notched, and dendritic complexity is not different from untreated controls in these groups. Tungsten particles, coated with the carbocyanine dye DiO, were delivered to cells in slices using the Helios Gene Gun System (BioRad) at 80 psi of helium gas pressure. Sections were then kept at 4° C for 24 hours on a shaker and post-fixed in 4% formalin for one hour. Sections were counterstained with DAPI (1:1000 in PBS) for 5 minutes, washed, and cover-slipped with ProLong Diamond Antifade Mountant, (Invitrogen Catalog # P36961). Slides were imaged at 10x by confocal microscopy (Zeiss) using Slidebook software. Images were captured within 3 days of DiO delivery to minimize dye diffusion. Cells were selected by morphology and cortical layer; only pyramidal-like cells of cortical layers IV-VI, adjacent to the mCherry band, with a prominent apical neurite, were collected. Most of the traced cells were from layer IV-V, but cells from layer VI were not excluded. In our samples, cells that strongly expressed mCherry were infrequently labeled with DiO, and therefore cells with readily detectable *mCherry* expression were generally not included in analysis.

An investigator blind to treatment traced cells, quantified total dendritic length and total nodes, and performed Sholl analyses on labeled neurons. Dendritic length, nodes, and Sholl intersections for cells in the contralateral hemisphere were indistinguishable from untreated controls, indicating that AAV8-*hSyn-mCherry* does not alter the morphology of pyramidal cells within the contralateral hemisphere **(Supplemental Fig. 1)**. Therefore, data are represented as a ratio of ipsilateral/contralateral within animal, to avoid influences from inter-animal variability in baseline brain structure, or minor differences in DiO intensity across bead batches. All Sholl analyses and quantification of dendritic length were done with a 10 μm radius using the Fiji plugin Simple Neurite Tracer. Approximately 14-18 cells per animal were analyzed.

### 2.4 Strahler Order

Cortical neuron tracings generated for Sholl analysis of cells from AAV1-*pCAG-FLEX-EGFP*-injected animals were used to further quantify changes to dendritic morphology using Strahler Order (SO), similar to ^40^. For SO analysis, all neurite tips are labeled “1”, and moving toward the soma, when two segments of the same SO meet, the resulting bifurcation (node) is labeled “n+1”. When two segments of different SO meet, the node is labeled with the higher of the SO numbers. The Strahler Number (SN) of a given neuron is the highest SO that is reached by that neuron. All SO analysis was performed by investigators blinded to condition.

### 2.5 Electrophysiology

#### Slice preparation

Acute coronal cortical slices were prepared from 11 to 14-week-old C57BL/6J mice, with or without injection of AAV1-*pCAG-FLEX-EGFP* (no CRE). Animals were anesthetized via inhalation isoflurane, and brains were rapidly dissected out. Slices (350 μm) were prepared using a Vibratome Series 1000 (Technical Products International) in ice-cold sucrose slicing solution, containing (in mM): 240 sucrose, 2.5 KCl, 10 Na-HEPES, 10 glucose, 1 CaCl□, 4 MgCl□, and 0.2 ascorbic acid, pH 7.3, bubbled with 100% oxygen. Slices were allowed to recover for 1-3 hr at 23-30° C in an incubation chamber containing high Mg^2+^ ACSF (in mM: 124 NaCl, 2.5 KCl, 26 NaHCO□, 1.25 NaH□PO□, 10 Glucose, 1 CaCl□, and 3 MgCl□, equilibrated with 95% O_2_ / 5% CO_2_). For recording, slices were transferred to an open recording chamber, submerged and perfused at 2 ml/min with standard ACSF (in mM: 124 NaCl, 2.5 KCl, 26 NaHCO□, 1.25 NaH□PO□, 25 Glucose, 2 CaCl□, and 1.3 MgCl□, equilibrated with 95% O_2_ / 5% CO_2_) at room temperature (RT). Pharmacological agents were introduced by switching the perfusion reservoir.

#### Whole cell patch-clamp

Pipettes (4–8 M□) were filled with intracellular solution containing (in mM) 108 Cesium gluconate, 20 HEPES, 0.4 EGTA, 2.8 NaCl, 5 TEACl, 4 Mg-ATP, 0.3 Na-GTP, and 10 phosphocreatine, and 6.5 biocytin HCl, pH 7.3, 290 mOsm. Currents were detected using a Multiclamp 700 B amplifier (Axon Instruments), conditioned with a 2 kHz Bessel filter, and digitized at 5 or 10 kHz with a Digidata 1322A (Axon Instruments). Whole-cell input resistance and series resistance were monitored by hyperpolarizing voltage test pulses (−5 mV) delivered before and after record acquisitions. Cells were excluded from analysis if the holding current (*I*_hold_) exceeded −100 pA, or if Rs was > 30 MΩ or changed by >30% during the recording.

mEPSC were recorded for at least 5 minutes from L4/5 pyramidal neurons of primary somatosensory cortex in the presence of 1 μm TTX (Hello Bio) and 100 μm Picrotoxin (Tocris). Data was analyzed offline using MiniAnalysis software (Synaptosoft, NJ, USA) and Origin 6.0 (Microcal Software, Inc.). The detection threshold was set at 6 pA. A total 300 events were analyzed for each cell. For cumulative probability plots, 200 consecutive events were sorted.

Spontaneous AMPA receptor-mediated EPSCs were continuously recorded for 15 minutes in the presence of 50 μm DL-APV (Tocris) and 100 μm Picrotoxin. To block calcium-permeable AMPA receptors in some experiments, NASPM trihydrochloride (Tocris) was introduced into the recording chamber (final bath concentration, 100 μm) after 5 min of stable recording using a 100 μl micropipette. 10 mM QX-314 (Sigma) was added into the intracellular solution to block Na^+^ channels. The inhibition of AMPA receptor sEPSC amplitude was calculated by comparing sEPSCs recorded during the first 5 min (prior to addition of NASPM) and from 5-10 min after addition of NASPM.

### 2.6 RNA Extraction and cDNA Synthesis

For mRNA analysis, performed 4 days after viral injection, mice were anesthetized with 5% isoflurane (inhaled) and brains were immediately dissected for RNA extraction. Injected and non-injected brain regions were excised and flash frozen in dry ice. RNA was extracted from desired brain regions using the Promega SV RNA isolation kit (Promega Catalog # Z3100), following manufacturer instructions. Briefly, the tissue was homogenized in SV RNA lysis buffer, SV RNA dilution buffer was added, and the sample was vortexed and centrifuged at 15,000 rpm for ten minutes. The supernatant was transferred to 95% EtOH, mixed and transferred to a spin column assembly. This assembly was centrifuged for one minute and the flow through was discarded. DNAse and then MnCl_2_ was added to yellow core buffer. This solution was added to the pellet, and the sample was incubated for 15 minutes at RT. SV DNAse stop solution was added, the sample was centrifuged for one minute, and the flow through was discarded. The sample was then washed in SV RNA wash solution and centrifuged twice. Flow through was again discarded and the spin basket was transferred to an elution tube. The sample was left standing for five minutes at RT and centrifuged for one minute. RNA samples were aliquoted and immediately frozen at −80° C. An aliquot was used to check for proper RNA extraction and quality by electrophoresis and nanodrop (Thermo Fisher Scientific Catalog # ND-2000).

For cDNA synthesis, sample RNA, 5x First Strand Buffer (Fisher Scientific Catalog # 18064022) and DNAse were combined and incubated at 37° C for one hour. The sample was heated to 65° C for 15 minutes, Anchored Oligo (dTvN)_22_ primer was added, and the sample was cooled to 37° C for twenty minutes. dNTP (Fisher Scientific Catalog # A25741), RNAsin (Fisher Scientific Catalog # PRN2511) and SuperScriptII-RT (Fisher Scientific Catalog # 18064022) were added, and the sample was incubated at 37° C for one hour, then stored at 4° C until needed.

### 2.7 Primer Design and RT-qPCR

For RT-qPCR, *H2-K* and *H2-D* primers were custom made using the IDT primer quest tool. *Iba-1, C3* and *Fyn* primers for qPCR were taken from Qvartskhava ^65^, Riihila ^66^ and Lim et al. ^67^, respectively. After viral infection, Kuchipudi et al. ^68^ found that *18s rRNA* was the least affected among tested genes, and therefore we used *18s rRNA* as a reference gene for this study. Primer sequences can be found in **Supplemental Table 1**.

A Master Mix of SYBR green (Fisher Scientific Catalog # A25741), GoTaq polymerase (Fisher Scientific Catalog # PRM3001) and RNA-free H_2_O (Thermo-Fisher Catalog # 10977-015) was made in advance and added to centrifuge tubes for each brain region. Two microliters of cDNA per region was added to their respective tubes. Master Mix plus cDNA was added to each well in a 96 well plate, followed by 1μl of primer mix containing both forward and reverse primers. The RT-qPCR program was then run in an Applied Biosystems Quant Studio3 System for 50 cycles. Melt curves were calculated to establish the quality of the data, and any flagged data was omitted. There were no detectable *mCherry* mRNA in sham surgery and untreated samples, so the lowest possible C_T_ values (50) were assigned to these samples. Data were exported to Excel for analysis and analyzed using standard methods ^69^. When referring to the gene, the name is italicized, and when referring to the protein, the name is not italicized.

### 2.8 Western Blotting and Cytokine Arrays

For protein analysis, mice were lightly anesthetized by 5% inhalation isoflurane and immediately decapitated for brain extraction three weeks post-injection. Motor and somatosensory cortex, hippocampus and cerebellum of both hemispheres were excised. Samples were homogenized, and protein concentration was determined by BCA analysis (20 μg of protein per lane) (Thermo-Fisher Catalog # 23225). Samples were diluted in sample buffer and boiled for 1 minute. SDS-PAGE electrophoresis was run at 100 V for 1 hour. Proteins were transferred from the gel to a PVDF membrane (Millipore Sigma Catalog # IPVH00010, BioRad Catalog # 1620174) at 350 mA for 30 minutes to 1 hour. Strips were cut and blocked in 5% BSA in TBST (80.6 g Na^+^Cl^-^ with 24.23 g Tris Base and pH to 7.6 with HCl for 10x TBS; add 1 ml Tween 20 to a 1:10 dilution in H_2_O for 1x TBST) for 1 hour at RT. Primary antibodies were applied in 5% BSA blocking buffer at several dilutions to determine the optimal concentration for 18-24 hours at 4° C on an orbital shaker. Membranes were washed 3 times for 5 minutes in TBST, and secondary antibodies were applied in 5% BSA in TBST for 1.5 hours at RT. Membranes were washed 5 times for 5 minutes in TBST, then immersed in SuperSignal West Pico Substrate (Thermo-Fisher Catalog # 34080) for 5 minutes. Because the molecular masses of C3 and the reference protein were similar, the membrane was stripped (Millipore Sigma Catalog # 2504) and re-probed for the reference protein after labeling for C3. Black lines in **Fig. 2j; Fig. 3b; Supplemental Fig. 4b, f** and **Supplemental Fig. 5f** are for clarity to distinguish between lanes and conditions in a single blot. When images were cropped, white spaces are present between conditions in image. See **Supplemental Table 2** for list of antibodies. A mouse cytokine array panel A (R&D Systems Catalog # ARY006) was used to assess cytokine responses to AAV1-*pCAG-FLEX-EGFP* three weeks post-injection. Cytokines included in the array (including alternative names): CXCL13/BLC/BCA-1, IL-5, M-CSF, C5a, IL-6, CCL2/JE/MCP-1, G-CSF, IL-7, CCL12/MCP-5, GM-CSF, IL-10, CXCL9/MIG, CCL1/I-309, IL-12 p70, CCL3/MIP-1 alpha, CCL11/Eotaxin IL-13, CCL4/MIP-1 beta, ICAM-1, CXCL2/MIP-2, IFN-γ, IL-17, CCL5/RANTES, IL-1 alpha/IL-1F1, IL-23, CXCL12/SDF-1, IL-1 beta/IL-1F2, IL-27, CCL17/TARC, IL-1ra/IL-1F3, CXCL10/IP-10, TIMP-1, IL-2, CXCL11/I-TAC, TNF-α, IL-3, CXCL1/KC, TREM-1 and IL-4. Manufacturer’s instructions were followed, with 300 μg of total protein loaded into each well. All cytokine measures include two technical replicates on the same membrane.

For both western blotting and cytokine arrays, images were taken in an Optimax X-ray film processor (model # 1170-4-0000), and film was scanned for analysis using ImageJ. Multiple exposures were taken, and appropriate exposures were used to quantify densitometry within the linear range of assay sensitivity.

### 2.9 Statistics

For all data sets a Shapiro-Wilk test was done to ensure data was normally distributed, and a two-tailed Grubb’s test was used to identify outliers. Two tailed unpaired t-tests assuming equal variance were used for all comparisons. All technical replicates were averaged. A Kolmogorov-Smirnov test was used for analysis of cumulative curves (*p< 0.01).

For qPCR: to adjust for differences in sample size and mRNA levels, relative expression (ΔC_T_) values for each gene were defined by subtracting the reference gene C_T_ value from the target gene’s C_T_ value. Values were then taken out of log by raising the C_T_ value by 2^(−CT)^ and multiplied by 100,000. To reduce type I errors arising from multiple comparisons, only virus-untreated, virus-sham and sham-untreated comparisons were performed. One-tailed t-tests were used to analyze *mCherry* mRNA levels, because changes in only one direction (rises) are possible from baseline.

